## Supplementary figures and images for "Construction of competing endogenous RNA interaction networks as prognostic markers in metastatic melanoma"

### Supplemental Figure 1

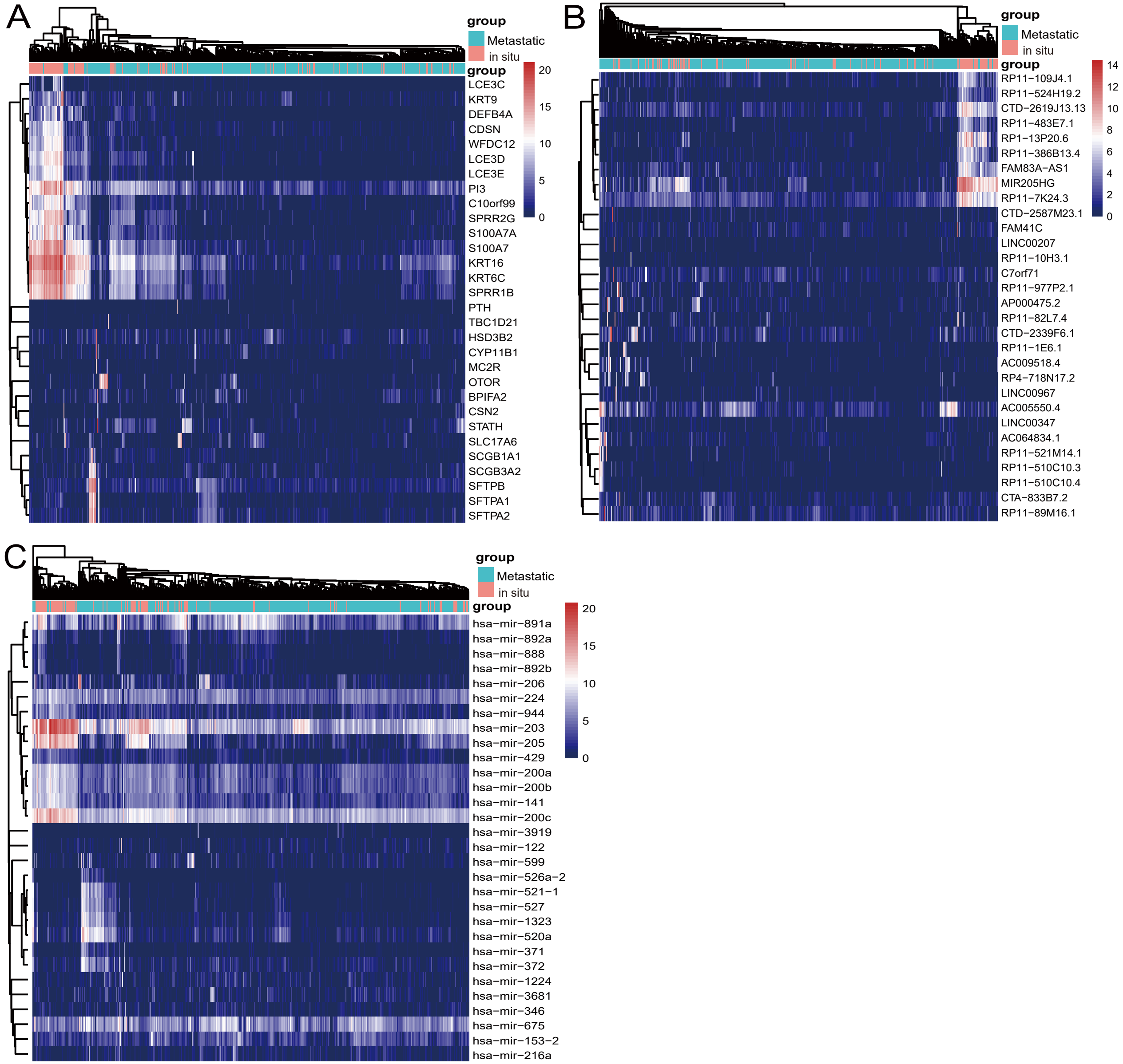

### Supplemental Figure 2

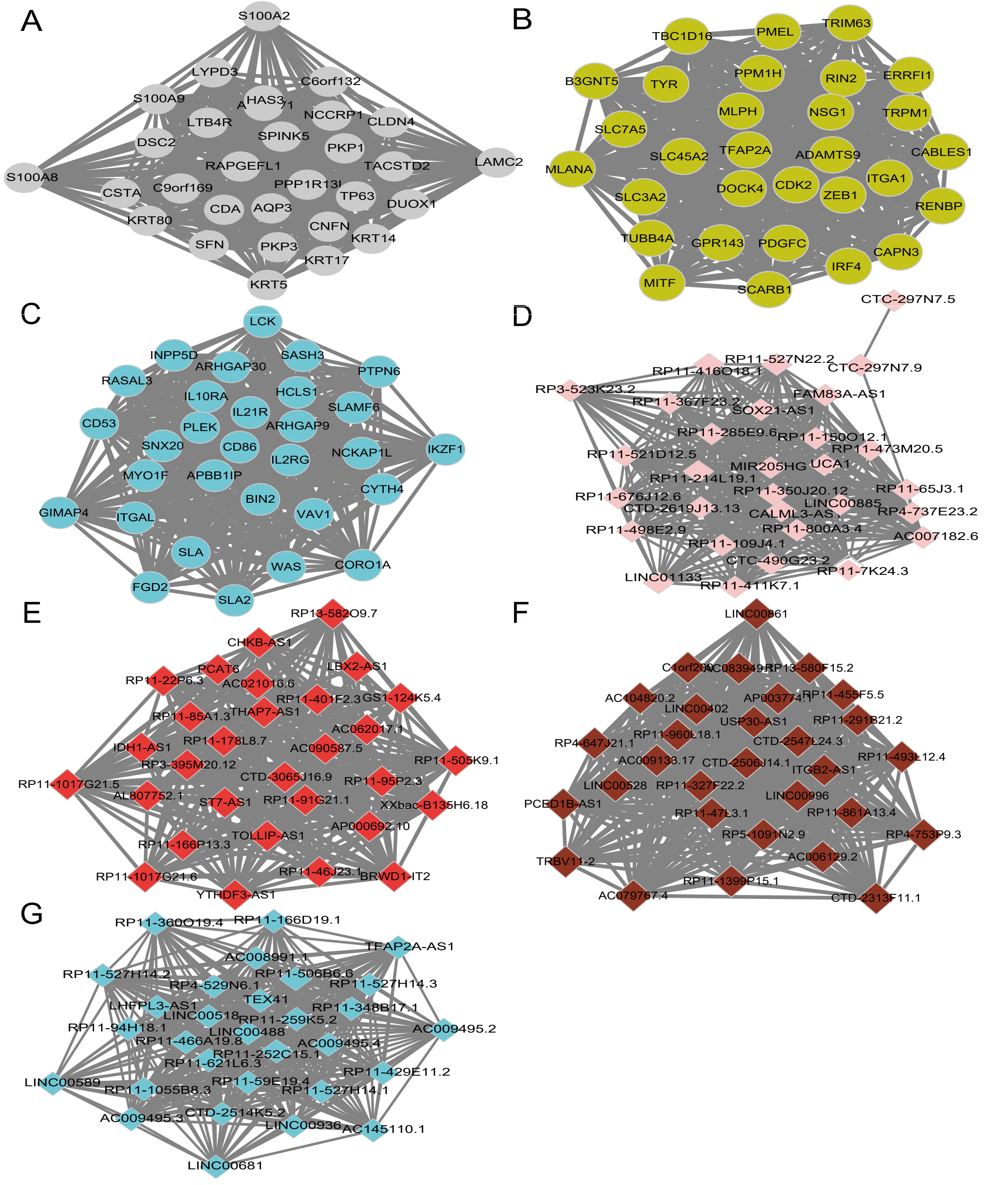

### Supplemental Figure 3

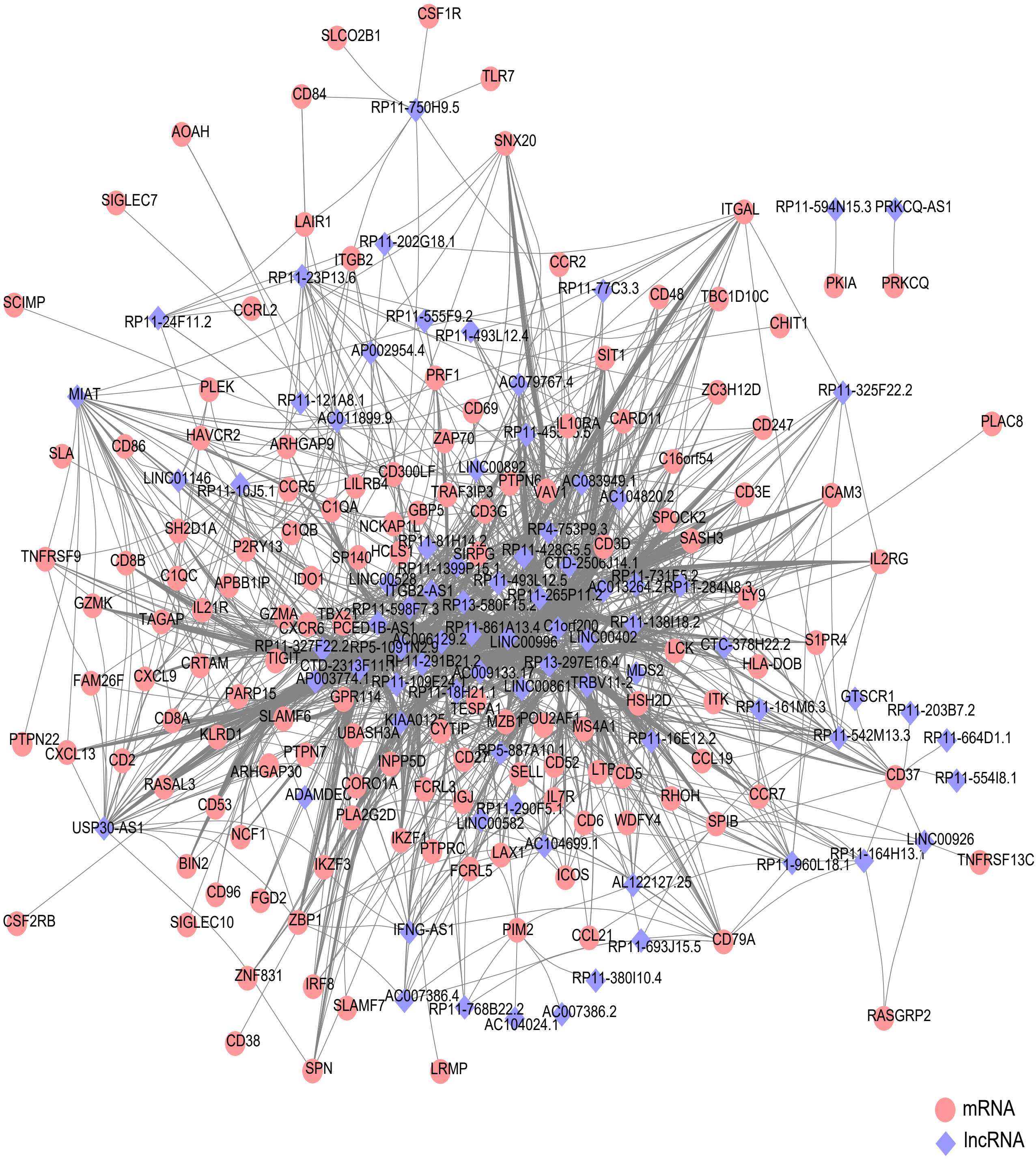
